# Genomic and Bioinformatic Insights into *Enterococcus faecalis* from Retail Meats in Nigeria

**DOI:** 10.1101/2025.04.15.648955

**Authors:** Charles Ayodeji Osunla, Ayorinde Akinbobola, Arif Elshafea, Esther Eyram Asare Yeboah, Olayemi Stephen Bakare, Aderonke Fayanju, Dorcas Oladayo Fatoba, Bright Boamah, Daniel Gyamfi Amoako

## Abstract

**Background:** *Enterococcus faecalis (E. faecalis)* is a commensal and opportunistic pathogen increasingly recognized for its antimicrobial resistance (AMR) and zoonotic potential. This study employs whole-genome sequencing (WGS) to characterize *E. faecalis* isolates from retail meat samples, focusing on antimicrobial resistance genes (ARGs), virulence determinants, mobile genetic elements, and phylogenomic relationships.

**Materials and Methods:** Fifty raw meat samples, including chicken (n=18), beef (n=17), and turkey (n=15), were collected from retail markets in Akungba-Akoko, Nigeria. *E. faecalis* isolates were identified using standard microbiological methods and subjected to antimicrobial susceptibility testing were further analysed using WGS.

**Results:** Ten *E. faecalis* isolates were recovered, with the highest prevalence in chicken (n=6), followed by beef (n=2) and turkey (n=2). All isolates were resistant to clindamycin, erythromycin, and tetracycline. Frequent ARGs included *aac(6’)-aph(2’’)*, *ant(6)-Ia*, *lsa(A)*, *erm(B)*, *tet(M)*, and *tet(L)*. Plasmid replicons *rep9c* and *repUS43* showed ST-specific associations with ST477 and ST16, respectively. MGEs such as *IS3*, *IS6*, *IS256*, and *IS1380* co-localized with ARGs and virulence determinants. Phylogenomic analysis revealed two major lineages, with ST477 distributed across meat types and ST16 restricted to chicken. Comparative genomic analysis with publicly available African *E. faecalis* isolates revealed distinct clonal lineages and geographic clustering across the continent.

**Conclusion:** The co-occurrence of multidrug resistance, virulence factors, and MGEs in foodborne *E. faecalis* poses a public health concern due to the risk of horizontal gene transfer and zoonotic spread. These findings underscore the need for genomic surveillance and antimicrobial stewardship in food systems, particularly in low- and middle-income countries.

## 1. Introduction

*Enterococcus faecalis* is a commensal bacterium of the gastrointestinal tracts of humans and animals, yet it has emerged as a notable opportunistic pathogen, especially in healthcare settings where multidrug-resistant (MDR) strains contribute to severe and difficult-to-treat infections (Farman et al., 2019). Beyond clinical contexts, its presence in food system particularly in raw meats raises significant concerns about its role in the dissemination of antimicrobial resistance genes (ARGs) and virulence factors via the food chain (de Mesquita Souza Saraiva et al., 2022). The species’ adaptability is bolstered by its remarkable capacity to acquire and transfer mobile genetic elements (MGEs), which facilitates horizontal gene transfer and complicates therapeutic strategies (Hegstad et al., 2010). These characteristics collectively pose a dual threat to both food safety and public health, necessitating a comprehensive understanding of its genomic architecture across diverse ecological niches.

Globally, genomic studies have examined the resistance mechanisms and genetic diversity of *E. faecalis* across clinical, livestock, and environmental settings (Daniel et al., 2017; Guan et al., 2024). However, substantial gaps remain in low-resource regions where genomic surveillance of foodborne isolates is limited (Okeke et al., 2022). In Nigeria, retail meat is a dietary staple, yet little is known about the genomic features of *E. faecalis* circulating in these products (Wada et al., 2020). Existing research has largely focused on phenotypic antibiotic resistance (Ndahi et al., 2023), with minimal exploration into the genetic determinants of resistance, virulence, and gene transfer that contribute to its pathogenic potential (Okeke et al., 2022). This is particularly concerning in a country where antibiotic use in agriculture remains poorly regulated and surveillance infrastructure is still evolving (Schnirring, 2023), potentially accelerating the emergence of MDR lineages.

The public health implications of foodborne *E. faecalis* are further underscored by increasing evidence of clonal transmission across animals, humans, and the environment (Monteiro Marques et al., 2023; Poulsen et al., 2012). Hospital-adapted lineages of *E. faecalis* have been shown to carry MGEs conferring resistance to clinically critical antibiotics such as vancomycin and β-lactams, as well as virulence genes that promote biofilm formation, immune evasion, and tissue invasion (Raven et al., 2016; Hourigan et al., 2024). If food-derived strains harbor similar genomic traits, this could signal a critical interface between agricultural and clinical reservoirs, a hypothesis that remains largely untested in key distribution hubs such as Akungba-Akoko, a prominent meat market in southwestern Nigeria (Alimi, 2013).

This study addresses this knowledge gap by performing a comprehensive genomic characterization of *E. faecalis* isolates recovered from retail meat in Akungba-Akoko. Utilizing whole-genome sequencing (WGS) and bioinformatics approaches, we aim to (1) assess the prevalence and diversity of ARGs, including those conferring resistance to critically important antimicrobials; (2) characterize virulence determinants associated with adhesion, biofilm formation, and immune evasion; and (3) investigate the mobile genetic elements (MGEs) facilitating gene exchange. These findings will contribute to our understanding of the genomic plasticity of *E. faecalis* in Nigeria’s food systems and inform mitigation strategies to reduce the public health risks posed by this emerging foodborne pathogen.

## 2. Materials and methods

### 2.1. Sample collection and study site

The study was carried out over a three-month period between April and June 2022: 50 samples of raw retail meat, including chicken (n=18), beef (n=17), and turkey (n=15), from the Ibaka market (7.473500° N, 5.736250° E) in Akungba Akoko, Nigeria. The Ibaka market is a rural periodic day market located in Akoko, which is the host community of Adekunle Ajasin University.

### 2.2. Isolation and identification of *Enterococcus faecalis* isolates

In the laboratory, each meat sample was aseptically homogenized. Smears of the homogenates were prepared and subjected to Gram staining to identify gram-positive cocci arranged in pairs or short chains, which are characteristic of *Enterococcus* species. For bacterial isolation, aliquots of the homogenized samples were inoculated onto blood agar plates and incubated aerobically at 37°C for 24–48 hours. Colonies exhibiting typical *Enterococcus* morphology were selected for further testing. Presumptive *Enterococcus* isolates were identified on the basis of their Gram staining characteristics and ability to grow on blood agar. Biochemical tests were performed to confirm that the isolates were *Enterococcus* species. These tests included the Voges‒Proskauer (VP) test for detecting acetoin production and the potassium tellurite (PT) test to assess the ability to reduce tellurite. Additionally, fermentation tests for glucose, lactose, and sucrose were conducted to evaluate the carbohydrate utilization profiles of the isolates. *Staphylococcus aureus* ATCC 29213 and *E. faecalis* ATCC 29212 served as negative and positive controls, respectively. Confirmed *E. faecalis* isolates were preserved by storing them in brain heart infusion broth (Difco) supplemented with 20% glycerol at −70°C for long-term storage.

### 2.3. Antibiotic susceptibility testing of *Enterococcus faecalis* strains

The antibiotic resistance profiles of *E. faecalis* isolates were determined via the disk diffusion method on Mueller‒Hinton agar (MHA), following the guidelines of the Clinical and Laboratory Standards Institute (CLSI, 2023). Overnight cultures of the isolates were used to prepare bacterial suspensions adjusted to a turbidity equivalent to a 0.5 McFarland standard. A sterile cotton swab was dipped into each suspension and evenly streaked across the entire surface of the MHA plates to ensure a uniform bacterial lawn. Commercial antibiotic disks (Hi-Media, India) were placed onto the inoculated plates via sterile forceps. The antibiotics used and their corresponding disk concentrations were as follows: tetracycline (30 μg), chloramphenicol (30 μg), streptomycin (10 μg), kanamycin (30 μg), erythromycin (15 μg), vancomycin (30 μg), clindamycin (2 μg), and tobramycin (10 μg). The plates were incubated aerobically at 37 °C for 18–24 hours. After incubation, the diameters of the inhibition zones around each antibiotic disk were measured in millimeters. *Staphylococcus aureus* ATCC 25923 was used as the control. The results were interpreted according to CLSI guidelines (CLSI, 2023), categorizing the isolates as susceptible, intermediate, or resistant to each antibiotic tested.

### 2.4. DNA Extraction, Whole-Genome Sequencing, and Assembly of *Enterococcus faecium* strains

Genomic DNA was extracted from *E. faecalis* isolates via the MasterPure™ Gram Positive DNA Purification Kit (Lucigen, Middleton, WI, USA) according to the manufacturer’s instructions. The quality and concentration of the extracted DNA were assessed via a NanoDrop 1000 spectrophotometer (Thermo Fisher Scientific, Waltham, MA, USA). The genomic DNA libraries were prepared via the Nextera XT DNA Library Preparation Kit (Illumina, San Diego, CA, USA) following the manufacturer’s protocol. Sequencing was performed on an Illumina NovaSeq 6000 system (Illumina, San Diego, CA, USA) to generate paired-end reads. The raw sequence reads were assembled into contigs via the Shovill pipeline version 1.0.4, which incorporates Trimmomatic version 0.38 for read trimming and quality control. Genome annotation was conducted via Prokka version 1.13.3.

### 2.5. Identification of resistance genes, virulence genes, plasmids and multi-locus sequence typing

Antimicrobial resistance genes were identified via ResFinder version 4.6.0 (http://genepi.food.dtu.dk/resfinder). Virulence genes were detected via VirulenceFinder version 2.0 (https://cge.food.dtu.dk/services/VirulenceFinder/). Plasmid types were determined by analysing the assembled genome sequences with PlasmidFinder version 2.1 (https://cge.food.dtu.dk/services/PlasmidFinder/). Multilocus sequence typing (MLST) was performed via the MLST tool version 2.0 (https://cge.food.dtu.dk/services/MLST/) to assign sequence types to the *E. faecalis* isolates.

### 2.6. Phylogenomic analysis and metadata insights

The de novo assembled contigs of the *E. faecalis* isolates were submitted to CSI Phylogeny version 1.4 (https://cge.cbs.dtu.dk/services/CSIPhylogeny-1.4), an online tool that identifies single nucleotide polymorphisms (SNPs) from whole-genome sequencing (WGS) data, filters and validates SNP positions, and infers phylogeny on the basis of concatenated SNP profiles. The *Enterococcus faecalis* ATCC BAA-2128 strain (Accession number: NAQY00000000.1) was used as an outgroup to root the tree, facilitating the assessment of phylogenetic relationships among the *E. faecalis* strains. The phylogenetic tree was visualized and annotated with isolate metadata, including demographic information, sequence types, resistome profiles, and mobile genetic elements (MGE), via ITOL version 7 (https://itol.embl.de). This approach provided a comprehensive analysis of the phylogenomic relationships among the isolates. Additionally, whole-genome sequences of *E. faecalis* isolates from Africa, curated from public databases such as PATRIC (https://www.patricbrc.org/) and NCBI between 2013 and 2023, were downloaded and included in the phylogenomic analysis to provide epidemiological and evolutionary context (Table S1). The trees were edited and visualized via FigTree version 1.4.4 (http://tree.bio.ed.ac.uk/software/figtree/). Isolates belonging to the same STs are highlighted with the same color, and isolates from the same geographical regions are labelled with the same text color to facilitate visual interpretation.

### 2.7. Nucleotide sequence

The sequences of the *E. faecalis* strains analysed in this study were deposited in the National Center for Biotechnology Information GenBank database under BioProject number **PRJNA928459**.

## 3. Results

### 3.1. Prevalence and Antibiotic Resistance Patterns of *E. faecalis* in Retail Meats

From the 50 retail meat samples analyzed, a total of 10 *Enterococcus faecalis* isolates were recovered, corresponding to an overall prevalence of 20%. The isolates were unevenly distributed among the meat types: chicken accounted for the highest number of isolates (n=6, 60%), followed by beef (n=2, 20%) and turkey (n=2, 20%). Antimicrobial susceptibility testing revealed consistent resistance profiles across the isolates. All strains exhibited complete resistance (100%) to clindamycin, erythromycin, and tetracycline, antibiotics commonly used in veterinary and clinical settings. Moderate resistance was observed against streptomycin (80%), and tobramycin (80%). In contrast, resistance to chloramphenicol was comparatively lower (20%), and no resistance to vancomycin was detected among any of the isolates. The distribution and co-occurrence of resistance phenotypes are illustrated in Figure 1 using an Upset plot.

**Figure 1.**
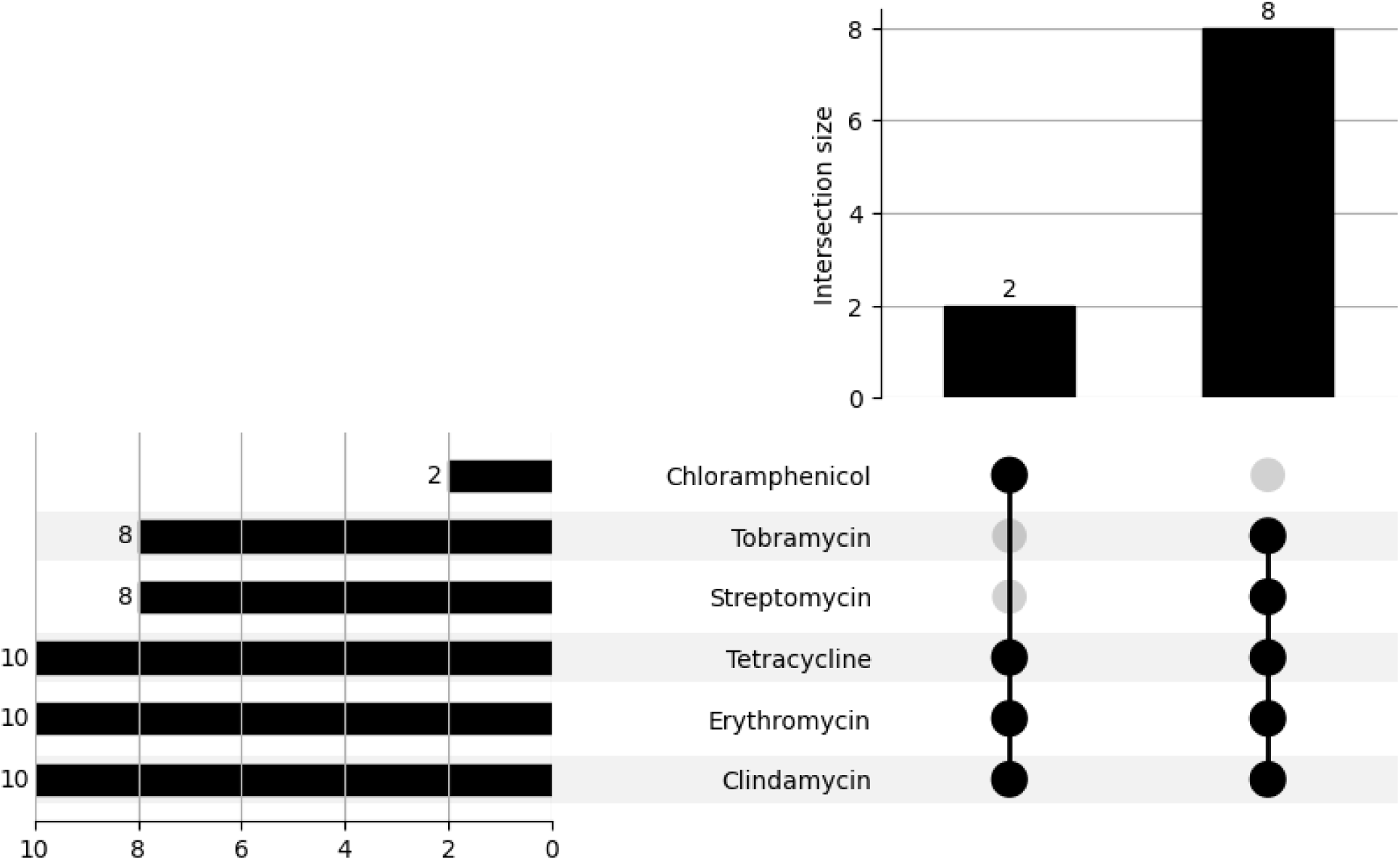
The Upset plot illustrates the distribution of antimicrobial resistance patterns across the isolates. The top bar chart quantifies unique combinations of antibiotic resistance among the isolates, while the horizontal bar chart on the left shows the number of isolates resistant to individual antibiotics. Co-resistance to clindamycin, erythromycin, tetracycline, and aminoglycosides (streptomycin and tobramycin) was common. Vancomycin was excluded from the visualization due to the absence of resistance. The arrangement highlights both dominant and infrequent co-resistance profiles across the dataset.

### 3.2. ARG profiles and mobile genetic elements

Genomic analysis identified a diverse repertoire of antimicrobial resistance genes (ARGs) among the *E. faecalis* isolates, many of which confer resistance to critically important antimicrobials. The most frequently detected ARGs included *aac(6’)-aph(2’’)* and *ant(6)-Ia* (aminoglycoside resistance), *lsa(A)* (lincosamide resistance), *erm(B)* (macrolide resistance), and *tet(M)* and *tet(L)* (tetracycline resistance). Notably, *dfrG* (trimethoprim resistance) and *catA8* (chloramphenicol resistance) were detected exclusively in two isolates recovered from retail chicken, suggesting lineage-specific acquisition or exposure to unique selective pressures.

Across meat sources, the core resistome was largely conserved; however, isolates NigeriaC1 and NigeriaC11 (from chicken) harbored a broader range of ARGs, reflecting potential differential antibiotic exposure in poultry production systems (Table 1). Analysis of mobile genetic elements (MGEs) revealed lineage- and source-specific patterns. The plasmid replicon *rep9c*was the most prevalent, detected in all isolates regardless of meat source and consistently associated with sequence type ST477. Conversely, *repUS43* was uniquely found in ST16 isolates from retail chicken, indicating a possible plasmid-lineage specificity.

**Table 1:**
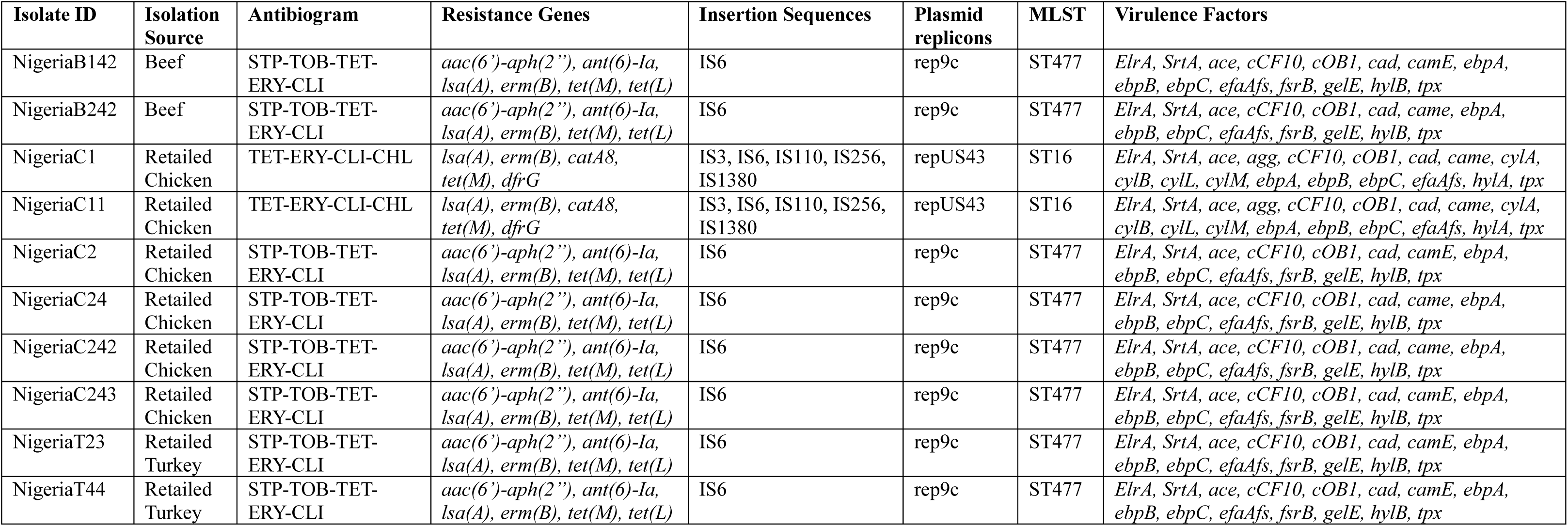
Genomic and Phenotypic Characteristics of *E. faecalis* Isolates.

Insertion sequences (ISs) played a prominent role in the resistome architecture. The IS6 family was frequently identified in ST477 isolates from all meat types. A unique combination of IS elements; *IS3*, *IS6*, *IS110*, *IS256*, and *IS1380* was observed only in ST16 isolates, co-localized with *dfrG* and *catA8*. Further analysis showed identified a resistance gene cassette comprising *aac(6’)-aph(2’’)*, *ant(6)-Ia*, and *tet(M)* embedded within a genomic region enriched with mobile genetic elements, including *IS1380* and *IS6* family transposases, as well as plasmid recombinase family proteins (Figure 2).

**Figure 2.**
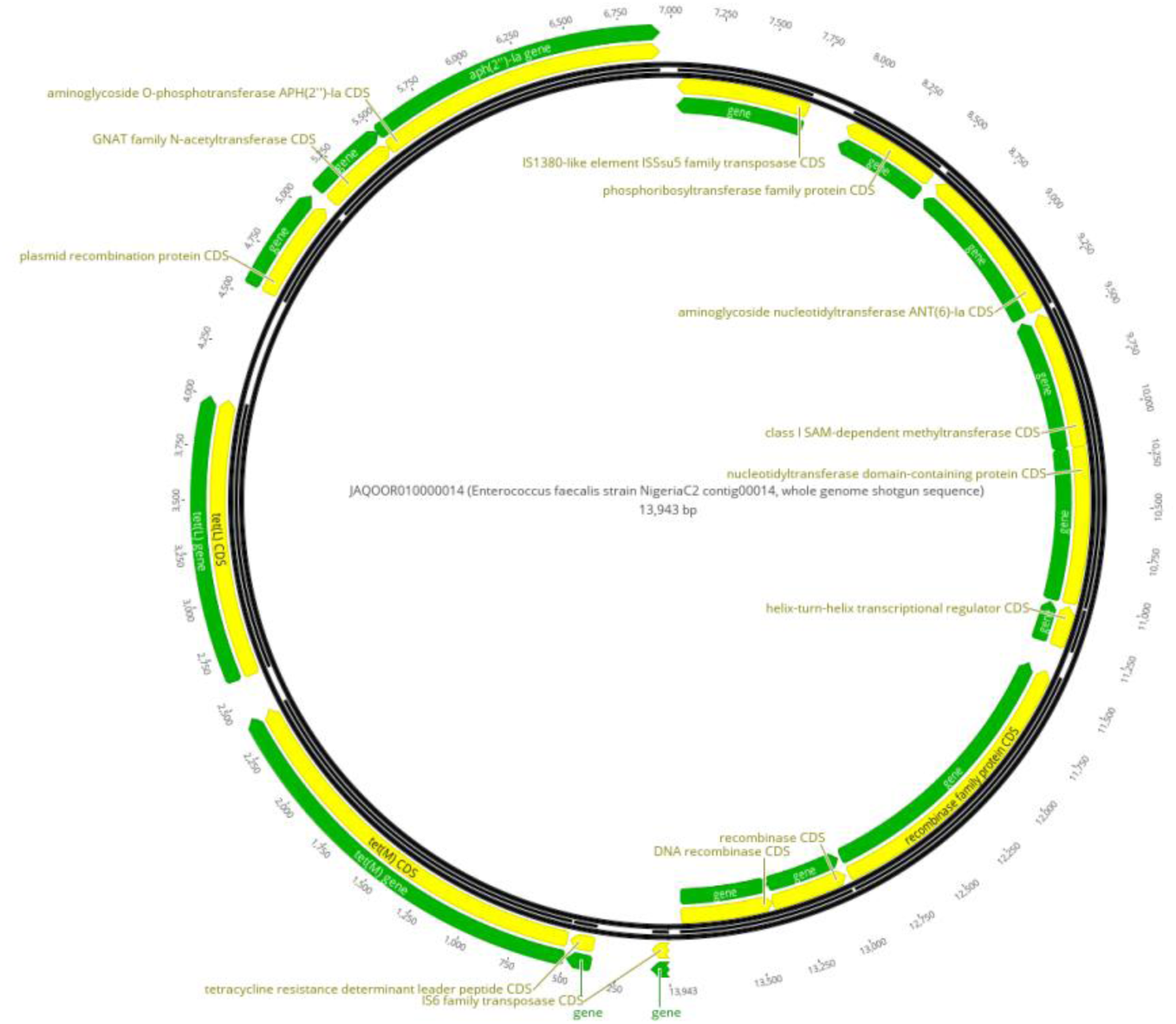
Circular genomic annotation of the genetic cassette carrying resistance genes in *Enterococcus faecalis* isolate NigeriaC2. This figure presents a visualization of the genomic region containing resistance determinants *aac(6’)-aph(2’’)*, *ant(6)-Ia*, and *tet(M)* in isolate NigeriaC2 (Accession number: JAQOOR010000014). The annotation highlights the relative positioning and orientation of these resistance genes alongside associated mobile genetic elements, including *IS1380* transposase, *IS6* family transposase, and plasmid recombinase family proteins. Genes are color-coded, with green representing resistance genes and regulatory elements, while yellow indicates protein-coding sequences (CDS). Arrows denote the transcriptional orientation of individual genes, providing insights into their synteny and potential functional interactions within the genomic context.

### 3.3. Virulence factors, sequence types and phylogenetic insights

Genome analysis revealed that *E. faecalis* isolates harbored an extensive repertoire of virulence factors associated with colonization, tissue invasion, and immune evasion. Conserved genes across all isolates included *ace* (collagen adhesion), *efaAfs* (endocarditis antigen), *ElrA* (adhesion), *SrtA* (anchoring protein), and the biofilm-associated pili genes *ebpA*, *ebpB*, and *ebpC*. Additionally, all isolates carried *cCF10* and *cOB1*, encoding aggregation substances that enhance horizontal gene transfer and biofilm development.

The distribution of virulence genes displayed clear sequence type (ST)-specific patterns. ST477, the most prevalent lineage, was recovered from beef, chicken, and turkey, and consistently carried *fsrB*, *gelE* (gelatinase), and *hylB* (hyaluronidase), key factors implicated in biofilm formation, extracellular matrix degradation, and immune modulation. In contrast, ST16 isolates (NigeriaC1 and NigeriaC11), found exclusively in chicken, exhibited a distinct virulence profile. These isolates harbored *agg*, *cylA*, *cylB*, *cylL*, and *cylM* genes encoding the cytolysin toxin complex, which contributes to tissue damage and enhanced pathogenicity. ST16 also possessed *hylA*, an alternative hyaluronidase variant, and *camE* (calcium-binding protein).

Plasmid replicons showed strong lineage associations. *rep9c* was universally detected in ST477 isolates and co-occurred with the *fsrB–gelE–hylB* virulence gene set. Conversely, *repUS43* was exclusive to ST16 and linked to the presence of cytolysin genes and *hylA*. Insertion sequences (ISs) were also associated with virulence profiles. ST477 isolates commonly carried *IS6*, while ST16 harbored a broader array of MGEs, including *IS3*, *IS110*, *IS256*, and *IS1380*, possibly facilitating mobilization of cytolysin and adhesion genes. Phylogenetic reconstruction using SNP-based analysis revealed two well-defined clades corresponding to ST16 and ST477 lineages (Figure 3). ST16 isolates clustered together and were uniquely associated with poultry, distinct ARGs, and virulence factors. ST477 formed a separate, more diverse clade encompassing isolates from all meat types and displaying a conserved resistome-virulome profile.

**Figure 3:**
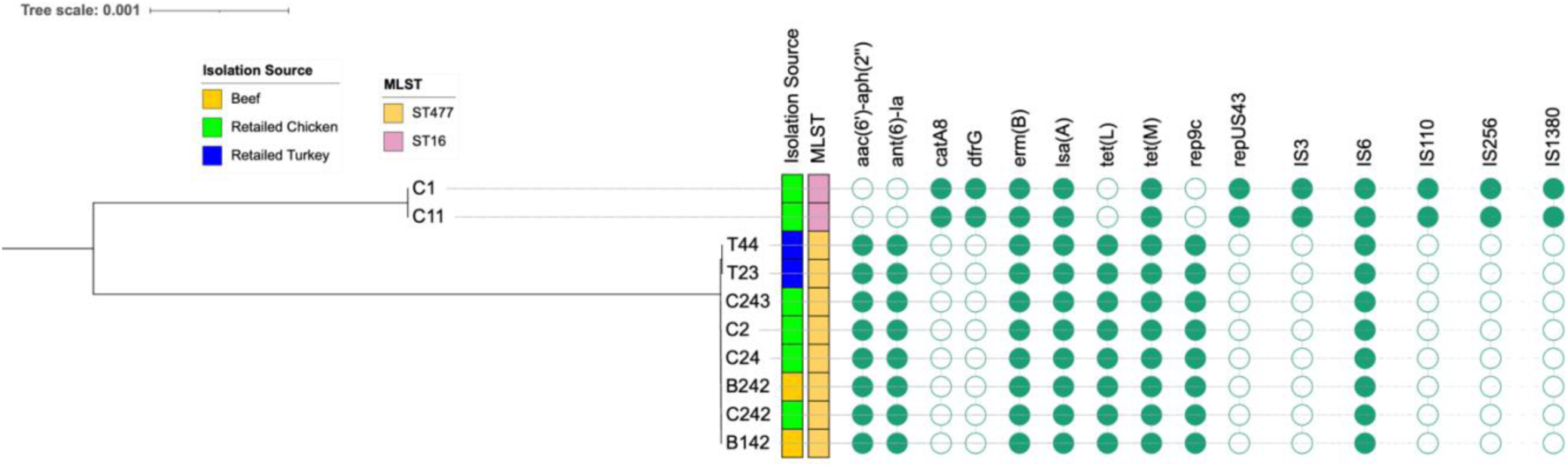
SNP-based maximum likelihood tree showing phylogenetic relationships among *E. faecium* isolates recovered from retail meat. The tree illustrates two main clades, each linked to specific sequence types (STs) and genomic characteristics. Annotations highlight key genomic features, including STs, isolation sources, resistance genes, plasmid replicons, and insertion sequences. Isolates are visually distinguished by color-coded boxes, indicating their distribution across different meat sources.

### 3.4. Comparative phylogenomic analysis and metadata insights of *E. faecalis* isolates from Africa

To contextualize the Nigerian *E. faecalis* isolates within broader regional dynamics, we conducted a comparative phylogenomic analysis of 149 publicly available African genomes collected between 2013 and 2023. South Africa contributed the highest number of isolates (n=60), followed by Tanzania (n=33) and Ghana (n=20), reflecting the uneven distribution of genomic surveillance efforts across the continent. Nigeria, the focus of this study, accounted for 10 isolates, while Cameroon, Zimbabwe, Tunisia, and Egypt contributed fewer (Figures 4 and 5a).

**Figure 4.**
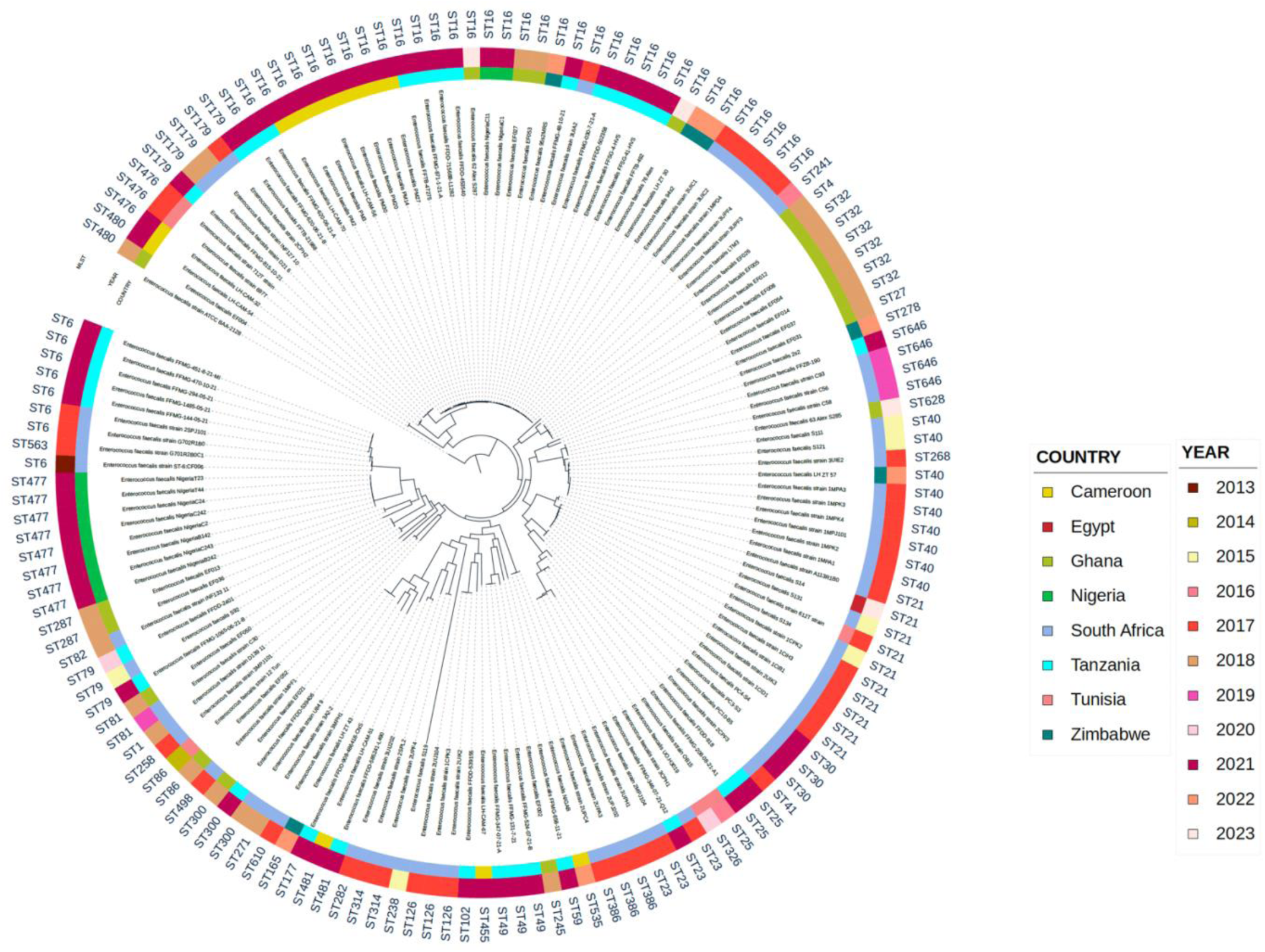
Maximum likelihood phylogenetic tree of *Enterococcus faecalis* isolates from African countries (2013–2023). A core-genome phylogenetic tree was constructed from 149 *E. faecalis* genomes retrieved from BV-BRC and annotated using iTOL. The tree is rooted using *E. faecalis* ATCC BAA-2128 (Accession: NAQY00000000.1) as the reference genome. The outer ring represents the sequence type (ST), the middle ring denotes the year of isolation, and the inner ring displays the country of origin. The phylogeny illustrates both temporal and geographic diversity of *E. faecalis* across the African continent.

**Figure 5.**
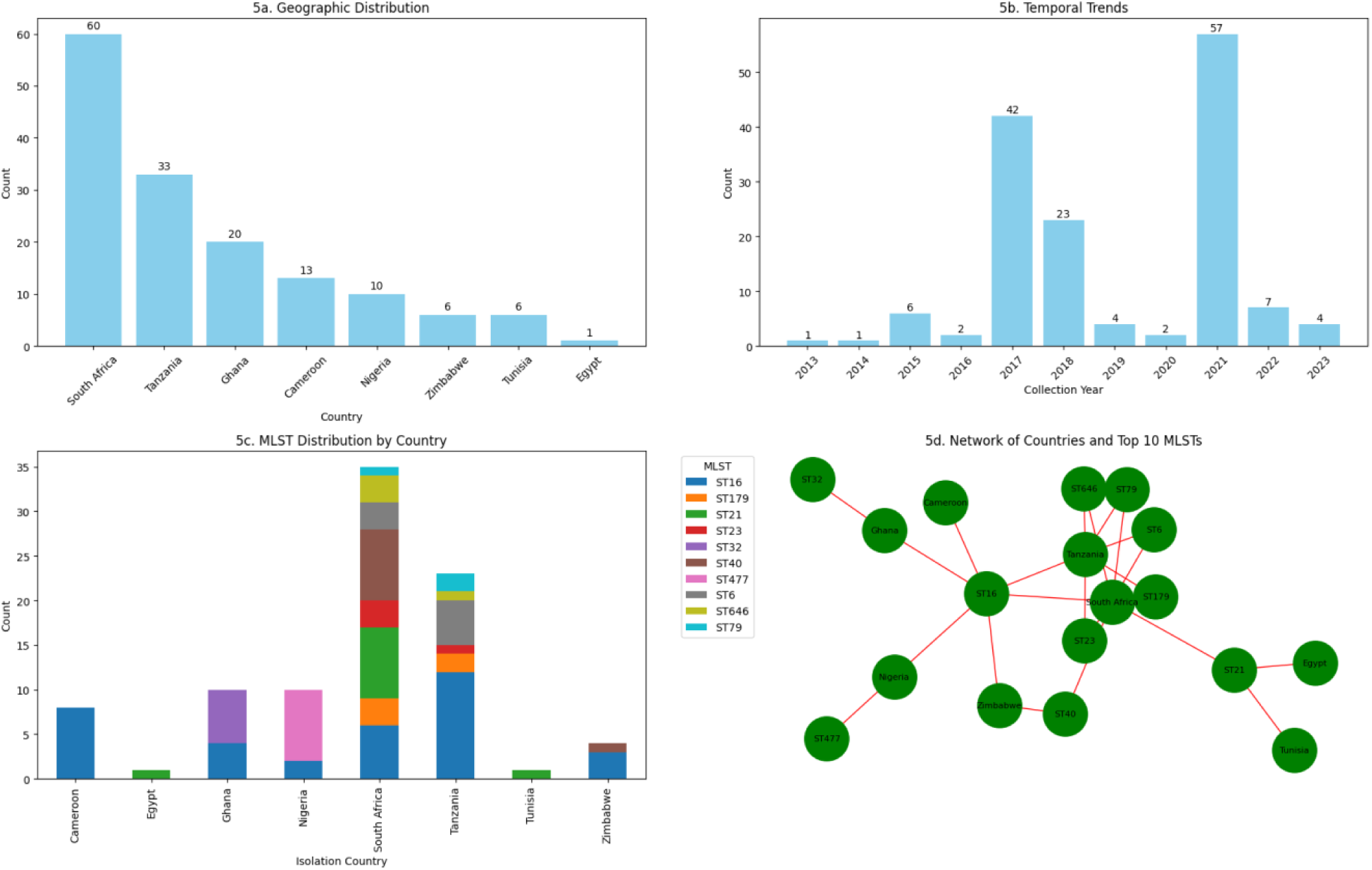
(a) Geographic distribution of *E. faecalis* isolates across Africa. A bar chart illustrating the number of *E. faecalis* isolates obtained from different African countries. (b) Temporal trends in *E. faecalis* isolations across Africa. A bar chart depicting the number of *E. faecalis* isolates collected per year from 2013 to 2023. (c) Distribution of major sequence types (*STs*) among African *E. faecalis* Isolates. A stacked bar chart showing the distribution of the most prevalent *STs* across different African countries. (d) A network graph visualizing relationships between African countries and their associated *E. faecalis STs*. Nodes represent countries and *STs*, with edges indicating connections based on isolate presence.

Temporally, most isolates were obtained between 2017 and 2021, with peaks in 2017 (n=42) and 2021 (n=57). Earlier years (2013–2016) were underrepresented, limiting historical comparisons but indicating a growing interest in enterococcal genomics in recent years (Figure 5b). Sequence type analysis identified 47 distinct STs across the dataset, with both shared and country-specific lineages. ST16 was the most widely distributed, detected in South Africa, Tanzania, Cameroon, and Ghana (Figure 5c). In contrast, ST477 found exclusively in the current Nigerian isolates appeared geographically restricted. Other notable country-specific lineages included ST21 (Tunisia and Egypt) and ST646 (South Africa). A network analysis of the top 10 STs and their country associations (Figure 5d) further illustrated these patterns. South Africa and Tanzania exhibited the highest ST diversity, with multiple connections to ST6, ST16, and ST646 (Figure 5d).

## 4. Discussion

The presence of *E. faecalis* in retail meats highlights the potential role of foodborne transmission in the dissemination of AMR (Conceição et al, 2023). Consistent with previous reports, chicken meat yielded the highest recovery rate of *E. faecalis*, reinforcing its role as a major reservoir for enterococci in food systems (Hayes et al., 2003; Aslam et al., 2012). Although not a classical foodborne pathogen, the species’ capacity to horizontally transfer resistance genes within the human gut microbiota elevates its public health significance (Krawczyk et al., 2021). These findings support calls for strengthened hygiene protocols and antimicrobial stewardship in poultry production, a sector characterized by intensive antimicrobial use (Conceição et al., 2023).

All isolates exhibited resistance to clindamycin, erythromycin, and tetracycline mirroring global resistance trends and suggesting sustained selection pressure from widespread use of these agents in animal husbandry (Arias & Murray, 2012; Lebreton et al., 2014; Landers et al., 2012). The absence of vancomycin resistance is encouraging; however, the species’ genomic flexibility raises concerns about future acquisition through horizontal gene transfer (van Hal et al., 2016). The detection of *catA8* among chicken isolates, despite the relatively low phenotypic resistance to chloramphenicol, signals the emergence of latent resistance and highlights the risk of resurgence (Bae et al., 2021). These findings reinforce the critical need for integrated AMR surveillance across food and clinical sectors, especially in high-burden regions (WHO, 2019).

Genomic analysis revealed a consistent resistome dominated by *aac(6’)-aph(2’’)*, *ant(6)- Ia*, *lsa(A)*, *erm(B)*, *tet(M)*, and *tet(L)* conferring resistance to aminoglycosides, macrolides, lincosamides, and tetracyclines. These genes, conserved across meat types and lineages, mirror globally recognized resistance profiles and likely reflect co-selection pressures exerted by agricultural antibiotic use (Hegstad et al., 2010). Of particular note, *dfrG* and *catA8* were confined to ST16 isolates from chicken, suggesting lineage-specific resistance acquisition possibly driven by poultry-associated selective environments (Kristich et al., 2014). These resistance determinants were not randomly distributed but instead embedded within complex mobile genetic architectures. A multidrug resistance gene cassette co-localizing *aac(6’)-aph(2’’)*, *ant(6)-Ia*, and *tet(M)* was detected within a genomic region enriched with IS6 and IS1380 transposases and plasmid recombinase genes, forming a transferable module capable of en bloc dissemination of resistance traits. Such modularity enhances the potential for inter-species gene flow, especially within gut microbiota of exposed consumers (Hegstad et al., 2010).

Plasmid replicons further delineated lineage-specific resistance pathways. The widespread *rep9c* replicon, present in all ST477 isolates, was consistently co-located with the *fsrB– gelE–hylB* virulence cluster, suggesting clonal expansion and vertical maintenance of resistance– virulence hybrids (Willems et al., 2012). In contrast, *repUS43* was exclusive to ST16 isolates, co-occurring with *catA8* and *dfrG*, reinforcing its role in ST-specific resistance dissemination (van Schaik et al., 2010). The structural linkage between plasmid types, insertion sequences, and ARGs underscores the dynamic interplay of vertical inheritance and horizontal transfer in shaping the resistome of *E. faecalis*. Insertion sequences (IS3, IS6, IS110, IS256, and IS1380) were found at multiple resistance loci and appear to drive genomic fluidity by facilitating recombination and gene mobilization. Their lineage-specific distribution highlights ongoing adaptive evolution under antimicrobial pressure. These findings illustrate a highly structured yet flexible resistance landscape in foodborne *E. faecalis*, propelled by mobile elements and reinforced by selective agricultural practices.

The widespread detection of virulence genes among *E. faecalis* isolates from retail meats reveals a pathogenic potential far beyond commensal behavior. Conserved adhesion factors including *ace*, *efaAfs*, *ElrA*, *SrtA*, and the *ebp* operon (*ebpA*, *ebpB*, *ebpC*) were uniformly present, underscoring a baseline capacity for epithelial colonization, biofilm formation, and immune modulation (Nallapareddy et al., 2006; Șchiopu et al., 2023). The concurrent presence of conjugative factors (*cCF10*, *cOB1*) suggests an additional role in promoting gene exchange within host or environmental niches, facilitating co-selection and persistence of resistance–virulence traits.

Lineage-specific virulence signatures were particularly striking. ST477 isolates recovered from beef, chicken, and turkey harbored the *fsrB–gelE–hylB* cluster, a constellation of genes associated with quorum sensing, extracellular matrix degradation, and immune evasion, previously linked to device- and bloodstream-associated infections (Van Tyne et al., 2013; Johnson et al., 2024). This combination of biofilm-promoting and immunomodulatory functions positions ST477 as a high-risk foodborne lineage with potential for clinical crossover. In contrast, ST16 isolates exclusively from chicken exhibited an enhanced virulence profile characterized by cytolysin operon genes (*cylA*, *cylB*, *cylL*, *cylM*) and *hylA*, features associated with epithelial disruption and pro-inflammatory host responses (Zheng et al., 2017). Notably, these isolates also carried *agg*, a plasmid-borne aggregation substance linked to increased virulence and conjugation efficiency. The co-occurrence of *repUS43* with this virulence suite suggests a plasmid-mediated mechanism facilitating the emergence of hypervirulent clones.

The convergence of resistance and virulence within specific plasmid backgrounds and sequence types especially ST16 and ST477 raises substantial public health concerns. Mobile elements, including IS3, IS110, IS256, and IS1380, were frequently associated with virulence loci, further implicating horizontal gene transfer in the amplification of pathogenic potential. These findings parallel global reports of virulence-enriched *E. faecalis* STs and affirm their relevance in zoonotic transmission and foodborne disease (McBride et al., 2007; Fiore et al., 2019). In this context, foodborne *E. faecalis* strains function not only as reservoirs of resistance but also as potential vectors for invasive disease, particularly in immunocompromised hosts. Their emergence in the food chain paired with genomic signatures linked to hospital-adapted strains underscores the urgency of a One Health surveillance framework integrating food safety, environmental monitoring, and clinical microbiology.

Comparative phylogenomics of *E. faecalis* isolates across Africa uncovered a geographically structured but genetically diverse population. South Africa contributed the highest number of isolates an observation likely shaped by differences in surveillance capacity, sequencing infrastructure, and public health prioritization. The scarcity of isolates from earlier years (2013– 2016) and the sharp rise in submissions post-2017, interrupted briefly by the COVID-19 pandemic, reflect both historical data gaps and the growing momentum of genomics-based AMR monitoring on the continent (Baker et al., 2023; Kajumbula et al., 2024; Tegally et al., 2022).

Temporal and geographic analyses revealed that sequence type ST16 was the most broadly distributed, identified in multiple countries and across diverse ecological contexts, suggesting a well-adapted and potentially mobile lineage (Zaheer et al., 2020; Monteiro Marques et al., 2023). Its widespread detection aligns with prior observations of ST16’s capacity for environmental persistence and inter-host transmission. In contrast, ST477 was restricted to Nigeria, while ST21 was confined to Tunisia and Egypt, implying localized evolutionary trajectories shaped by selective pressures such as antimicrobial usage patterns, ecological boundaries, and food production systems (Baquero et al., 2021; Bottery et al., 2021). Network-based analysis reinforced these findings by revealing distinct ST-country associations, with South Africa and Tanzania exhibiting the highest sequence type diversity. These clusters suggest transboundary transmission possibly facilitated by trade, food import/export routes, or shared agricultural practices (Tatem et al., 2006; Salem et al., 2023). Conversely, the geographic confinement of ST477 to Nigeria raises concerns about the emergence of a potentially endemic, foodborne high-risk clone with a stable resistome and virulome signature.

These patterns support a dual model of *E. faecalis* evolution in Africa: one shaped by clonal expansion of regionally successful lineages, and another driven by horizontal gene transfer across environmental and host reservoirs. This genomic duality complicates control efforts and underscores the need for longitudinal, cross-sectoral surveillance. The convergence of clinically relevant traits in isolates recovered from food reinforces the risk of zoonotic spill-over, particularly in settings with limited food safety regulation and AMR control. In light of these findings, a coordinated genomic surveillance strategy that integrates human, animal, and environmental health guided by the One Health framework is essential for tracking the emergence, evolution, and dissemination of high-risk *E. faecalis* clones in Africa. This study, while limited by its modest sample size and geographic scope, contributes a critical dataset to the continental AMR landscape and provides a foundation for future longitudinal studies on *E. faecalis* as a foodborne pathogen of increasing public health concern.

## Conclusion

This study provides critical genomic insights into the antimicrobial resistance and virulence landscape of *Enterococcus faecalis* isolates recovered from retail meats in Nigeria, underscoring the role of food systems as reservoirs and potential amplifiers of clinically relevant pathogens. The identification of high-risk lineages such as ST477 and ST16 harboring multidrug resistance determinants, virulence genes, and mobile genetic elements highlights the convergence of resistance and pathogenicity within the food chain. The presence of plasmid-encoded resistance–virulence modules and the widespread occurrence of insertion sequences suggest an active mobilome facilitating gene exchange and adaptation across ecological boundaries. Comparative phylogenomic analysis across Africa revealed geographically structured transmission dynamics, marked by the emergence of regionally dominant clones and country-specific evolutionary trajectories. These findings reflect broader challenges in AMR control, particularly in low- and middle-income countries where food safety infrastructure and genomic surveillance remain limited.

To mitigate the growing threat of foodborne antimicrobial resistance, we advocate for enhanced genomic monitoring of foodborne pathogens, stringent regulation of antimicrobial use in agriculture, and integration of One Health strategies across the human–animal–environment interface. The insights presented here serve as a foundation for future regional surveillance initiatives and emphasize the need for proactive, genomics-informed interventions to safeguard public health.

## Declarations

### Ethics Approval

Institutional approval for the study protocol and sampling approach was granted by the Department of Microbiology, Adekunle Ajasin University (Approval Reference: DM:AAU/2021). All meat samples were obtained from open retail markets in Akungba-Akoko, Nigeria, and were purchased anonymously as part of routine food supply, without involving live animals or interventions. The objectives of the study were clearly explained to meat vendors to ensure transparency and voluntary participation in the sampling process.

### Clinical Trial

Not applicable

## Data Availability

The sequence data that support the findings of this study has been deposited in GenBank and assigned accession numbers under BioProject **PRJNA928459**. All other data supporting this study findings are available within the manuscript and supplementary material.

